# Tryptophan-aspartic acid containing peptide analog from coronin 1 inhibits model membrane fusion and enveloped viral infection in cells

**DOI:** 10.1101/2025.05.14.653931

**Authors:** Swaratmika Pandia, Bushra Qazi, Vaishali Viswakarma, Shruti Gautam, Chetan D Meshram, Hirak Chakraborty, Sourav Haldar

## Abstract

Membrane fusion is a crucial step in the infection cycle of an enveloped virus, and the development of fusion inhibitors could lead to broad-spectrum antivirals beyond the one-bug-one-drug paradigm. In our continued effort to design peptide-based fusion inhibitors that block fusion by modulating membrane physical properties rather than targeting viral proteins, we have designed a tryptophan-aspartic acid (WD) containing peptide analog, mGG21, from coronin 1. Coronin 1 has been implicated in preventing the fusion of live mycobacteria containing phagosomes with lysosomes. The mGG-21 displays around 60% inhibition in fusion pore formation (complete fusion) in model membranes by increasing the acyl chain ordering of the membrane, regardless of the cholesterol content of the membrane, unlike previously designed predecessors with 20-30 % inhibition activity. Further, we show that mGG-21 inhibits Influenza and Chikungunya virus infection in cellular models without exerting any toxicity. Taken together, our findings underscore the importance of WD repeats in designing broad-range viral fusion inhibitors.

## Introduction

Membrane fusion is a dynamic process that enables the merging of two lipid bilayer-enclosed compartments, facilitating the transfer of content or cargo from one compartment to another. Membrane fusion plays a ubiquitous role in multiple cellular functions such as fertilization, cellular trafficking, neurotransmission, exo- and endocytosis.^1–3^ Enveloped viruses like human immunodeficiency virus (HIV), influenza, severe acute respiratory syndrome coronavirus (SARS-CoV), dengue, Chikungunya, and others utilize the same process to invade the host cells.^4–10^ Enveloped viruses either fuse directly with the plasma membranes of the host cells or with the endosomal membranes after endocytosis ^8, 11^ The fusion proteins present in the viral envelope catalyze the process by undergoing conformational transformations that bring two juxtaposed membranes close to establish molecular contact.^12, 13^ The insertion of the N-terminal peptide (fusion peptide of fusion loop) of the fusion protein further modulates the properties of the host cell membranes to facilitate virus-endosome (or virus-plasma membrane) fusion, which permits the transfer of the viral genome to the host cell.^14^ Formation of a six-helix bundle (6HB) through the interaction between N- and C-heptad regions of the fusion protein is a hallmark of its fusion-active conformation.^15–17^ A closer look at the viral fusion processes reveals that two bilayers merge to form a single bilayer through multiple non-bilayer intermediates, and the intermixing of the contents allows the transfer of the viral genetic material to the host cells.^18–22^ Therefore, viral fusion inhibitors represent a promising antiviral strategy. Several efforts have been made to block the fusion between the viral envelope and the host cell membranes to combat the infection process.^23–25^ Peptide-based viral fusion inhibitors represent potential antiviral drugs.^26^ One of the common strategies is to inhibit the formation of 6HB by preoccupying the C- or N-heptad region. Peptides similar to the sequence of the C-heptad region of a particular fusion protein bind to the N-heptad region and vice versa, and block the formation of 6HB, thereby inhibiting the fusion.^27–29^ The C-heptad, sequence of the spike protein from SARS-CoV, effectively inhibits viral entry by binding to the N-heptad region *in vitro*. However, its antiviral efficacy is low *in vivo*. Subsequent reports demonstrated that a lipid-conjugated C-heptad peptide displayed better inhibitory efficiency.^30, 31^ The membrane-proximal external region (MPER) sequence of the SARS-CoV-2 spike protein was further demonstrated as an ideal template for the design of fusion-inhibitory peptides.^32^ However, this approach is only viable for the viruses that fuse at the plasma membranes. T1249, Cp32M, T20, T2635, and Sifuvirtide are some examples of C-peptides, and NCCG-gp41, IQN17, and IQN23 are some examples of N-peptides that inhibit the entry of HIV.^33^ T20 (also known as Enfuvirtide or Fuzen), a 36-amino acid residue peptide derived from the C-heptad region of gp41, is the first peptide-based fusion inhibitor designed to prevent HIV entry.^34, 35^ Other peptide-based inhibitors do not block the 6HB formation but incapacitate the fusion protein to induce fusion. DN59 from dengue and WN83 from West Nile virus, with positive Wimley-White Interfacial Hydrophobicity Scale, proved to be most effective against dengue and West Nile virus infection, respectively, with IC_50_ values around 10 µM.^36^ On the other hand, several small molecules, such as cepharathine, abemaciclib, umifenovir, and osimertinib displayed decent entry inhibition against SARS-CoV infection.^30, 37^

However, these inhibitors target the fusion protein of a particular virus and hence are specific against a certain virus. Moreover, they might fail to deal with the mutations in the domain where they explicitly bind. Because of their specificity against a particular virus, this approach of inhibitor development is traditionally known as the ‘one-bug-one-drug’ paradigm.^38^ Therefore, these inhibitors are not adequate to address the continuously emerging and remerging viral diseases and raising the urge to develop broad-spectrum inhibitors that can work against the entry of multiple viruses. We have successfully exploited the WD-containing peptides from coronin 1, a phagosomal coat protein, to design inhibitor peptides that inhibit model membrane fusion and viral infection in cells.^39–43^ Coronin 1 plays a key role in inhibiting lysosomal fusion with the mycobacterium-containing phagosome.^44^ Despite being a phagosomal protein, coronin 1 is recruited to the phagosomal surface when the phagosome contains live mycobacteria.^45, 46^ The WD repeats draw special attention in designing inhibitor peptides as the position of these repeats is conserved across species.^47^ Even the lipidated WD dipeptide and lipidated branched D(WD)_2_ peptide inhibit the entry of influenza and murine coronaviruses.^48, 49^ We have earlier demonstrated that a 21 amino acid sequence from coronin 1, GCDNVIMV**WD**VGTGAAMLTLG (GG-21), displays about 25% inhibition during the fusion of small unilamellar vesicles (SUVs) containing varying amounts of membrane cholesterol.^42^ Based on the GG-21 sequence, we have designed a novel peptide, **mGG-21,** with two WD repeats close to N- and C-terminals (mGG-21: GC**WD**VILVAVVGTGAAV**WD**LG) to increase the fusion inhibitory efficiency. Our findings indicate that the mGG-21 blocks approximately 60% of fusion when tested in the SUV-SUV fusion assay. Measurement of depth-dependent membrane organization and dynamics utilizing fluorescence spectroscopy shows that the peptide inhibits fusion, probably by ordering the acyl chain region of the membrane. Since fusion is a crucial step in the viral infection cycle, we envisaged that mGG-21 can prevent viral infection in cells, conceivably by inhibiting viral fusion. To test this hypothesis, we have investigated the antiviral activity of mGG-21 against two different enveloped viruses, Influenza (with class I fusion protein) and Chikungunya (with class II fusion protein). Utilizing a range of biochemical and virological methods, we demonstrate that mGG-21 inhibits Influenza and chikungunya virus infection in cellular models without exerting any cellular toxicity. Our findings suggest a novel approach to designing a broad-spectrum antiviral strategy.

## Materials and Methods

Details of materials and methods have been included in the Supporting Information.

## Results

### Peptide Design and Hydrophobicity Profile

In this work, we have designed a novel coronin 1-derived peptide, mGG-21, by mutating the sequence of the GG-21 peptide. Both the GG-21 and mGG-21 are hydrophobic peptides, nonetheless, the hydrophobicity of mGG-21 is partially higher than GG-21 (**Fig. 1**). A peptide with higher hydrophobicity is expected to enhance its membrane partitioning ability. Two methionine residues of GG-21 have been substituted with leucine (7^th^ position) and valine (17^th^ position) to make the mGG-21 more hydrophobic. The residue-wise average hydrophobicity values were calculated by taking a running average of 5 amino acids using the expasy ProtScale algorithm.

**Figure 1.**
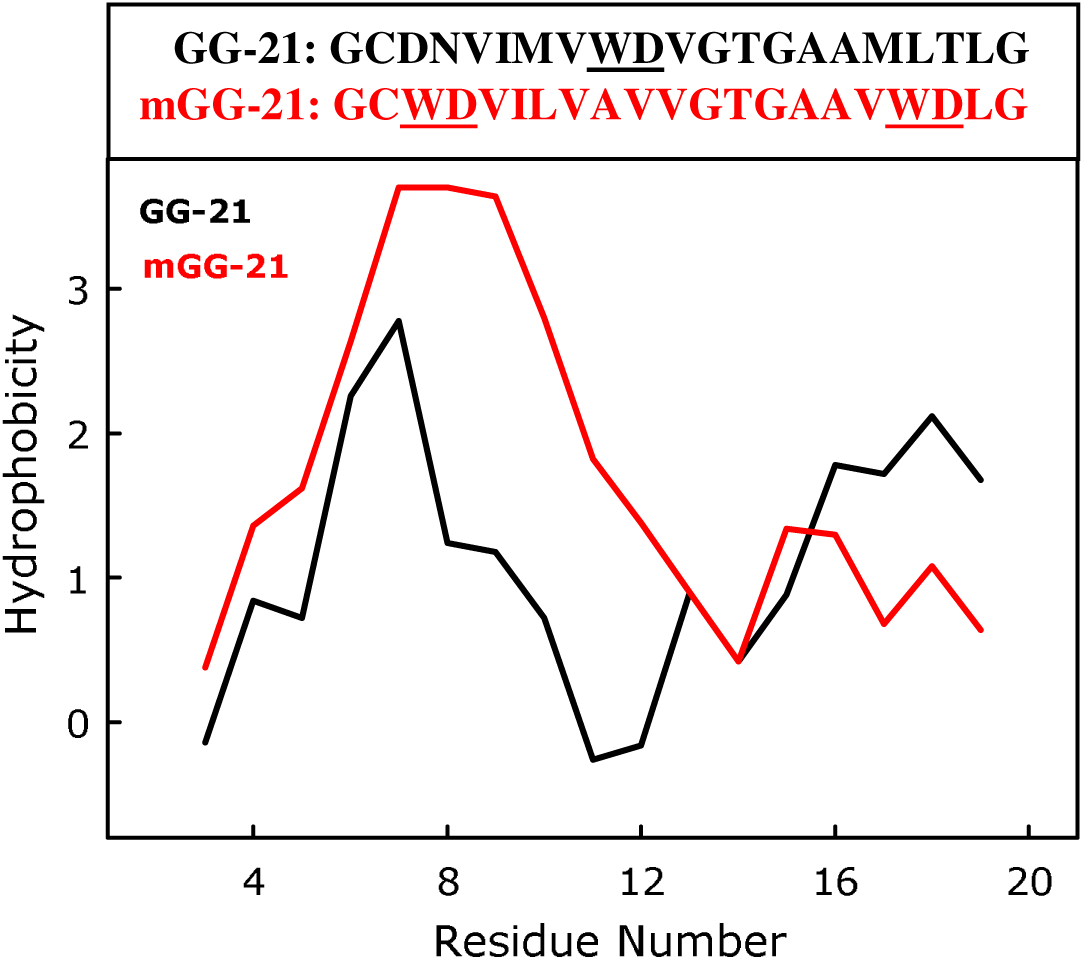
Plot of residue-wise hydrophobicity profile of GG-21 (black) and mGG-21 (red). The hydrophobicity was calculated using the Expasy-ProtScale algorithm with a window size of 5. The WDs in the peptide sequence are shown in bold and underlined.

### Partitioning of the mGG-21 peptide in membranes

The binding or partitioning of mGG-21 peptide to membranes with DOPC/DOPE/DOPG (60/30/10 mol%), DOPC/DOPE/DOPG/CH (50/30/10/10 mol%), and DOPC/DOPE/DOPG/CH (40/30/10/20 mol%) was evaluated by monitoring the fluorescence spectra of tryptophan present in the peptide in the absence and the presence of membranes. The tryptophan was excited at 295 nm, and the emission spectra were measured from 310 to 420 nm. The tryptophan fluorescence intensity increases significantly in the presence of membranes with various lipid compositions, with a concomitant blue shift in the fluorescence spectra (**Fig. 2**). The maximum emission of tryptophan shifts from 348 nm to 345 nm in the presence of the membrane. Both the enhancement of fluorescence intensity and blue shift of emission maximum imply the membrane partitioning of the peptide as tryptophan displays properties of a relatively hydrophobic environment in the presence of the membrane. However, the fluorescence intensity does not vary much with the varying compositions of the membrane, indicating a similar environment of tryptophan in membranes with various lipid compositions.

**Figure 2.**
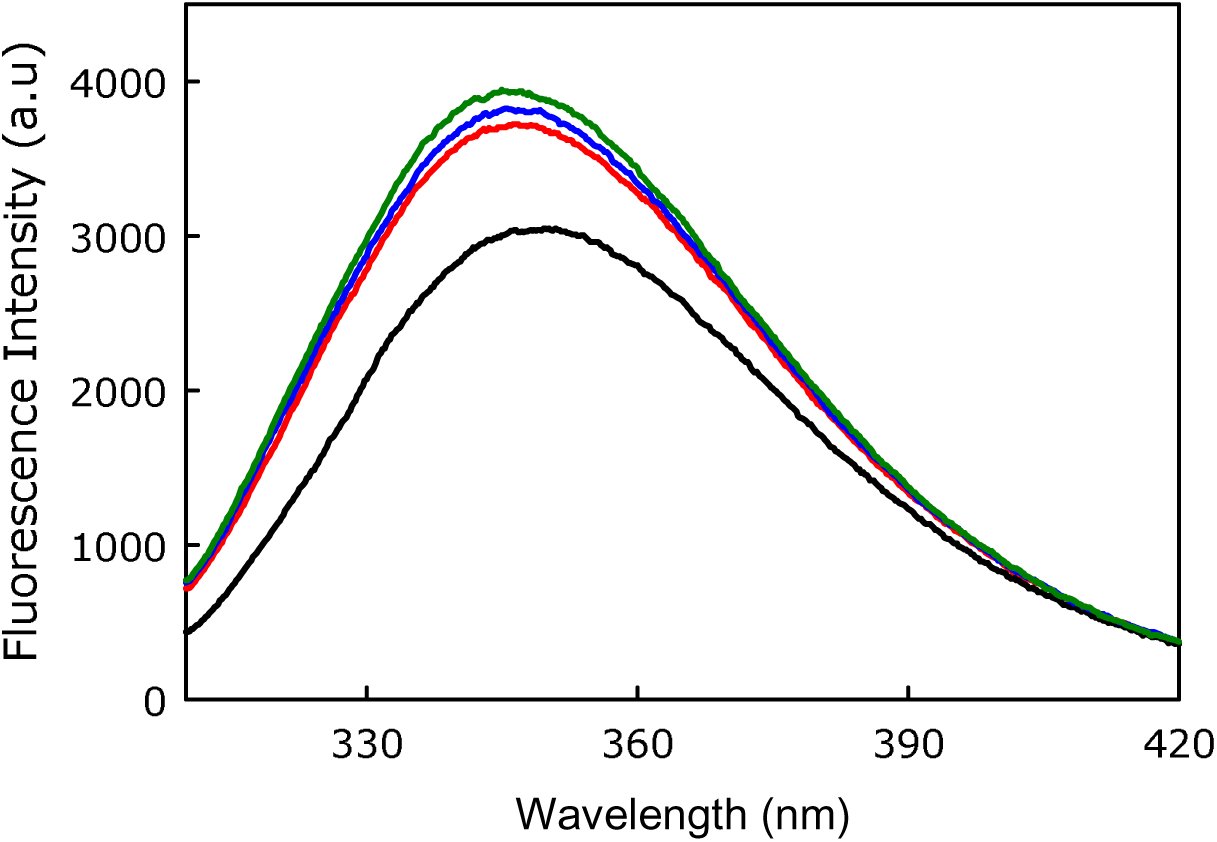
Fluorescence spectra of tryptophan present in mGG-21 in buffer (black), DOPC/DOPE/DOPG (60/30/10 mol%, red), DOPC/DOPE/DOPG/CH (50/30/10/10 mol%, blue), and DOPC/DOPE/DOPG/CH (40/30/10/20 mol%, green). Results are shown for a lipid-to-peptide ratio of 100:1 for measurements in the presence of membranes. Measurements were performed in 10 mM TES, 100 mM NaCl, 1 mM CaCl_2,_ and 1 mM EDTA, pH 7.4, at a total lipid concentration of 200 μM. The spectra are averages of at least three independent measurements.

### Effect of mGG-21 on polyethylene glycol (PEG)-mediated vesicle fusion

We have carried out PEG-mediated fusion between small unilamellar vesicles (SUVs) having varied lipid compositions to evaluate the fusion inhibitory effect of mGG-21. The fusion was induced by 6% (w/v) (PEG). It is well established that a small amount of PEG does not affect the physical properties of membranes, conformation, and the depth of penetration of the peptide into the membrane. The fusion observables, like lipid mixing (LM), content mixing (CM), and content leakage (L), were measured using the methodologies mentioned in the Materials and Methods section (Supporting Information). The PEG-induced fusion between SUVs having lipid compositions of DOPC/DOPE/DOPG (60/30/10 mol%), DOPC/DOPE/DOPG/CH (50/30/10/10 mol%), and DOPC/DOPE/DOPG/CH (40/30/10/20 mol%) was carried out both in the presence and absence of the peptide mGG-21. The vesicular fusion in the absence of peptide was considered as the control for all measurements, and the inhibitory efficacy of the peptide was examined against the respective control. The lipid-to-peptide concentration ratio was kept at 100:1 to maintain the physiological relevance of the low concentration of peptide. The percentage of LM, CM, and L was calculated using equations 2, 3, and 4, respectively (Materials and Methods section in Supporting Information). The kinetic data were analyzed using SigmaPlot (San Jose, CA) in a double exponential equation to determine the extent of three fusion observables at time infinity.

The kinetics of LM, CM, and L during the fusion of DOPC/DOPE/DOPG (60/30/10 mol%) membranes in the absence and presence of mGG-21 are displayed in **Fig. 3**(**A-C**). Our results demonstrate that the mGG-21 peptide promoted LM by nearly 26% while inhibiting CM by 59%. The peptide-induced reduction of L by 30% supports the rigidification of the membrane in the presence of mGG-21. The enhancement of LM with associated inhibition of CM suggests that mGG-21 promotes hemifusion formation and stabilizes it so that the fusion reaction does not proceed toward pore formation. Pore formation is considered the most essential step in membrane fusion as it involves the intermixing of contents.^50^ The viral genetic material cannot be transferred to the host cell if the pore formation is inhibited.

**Figure 3.**
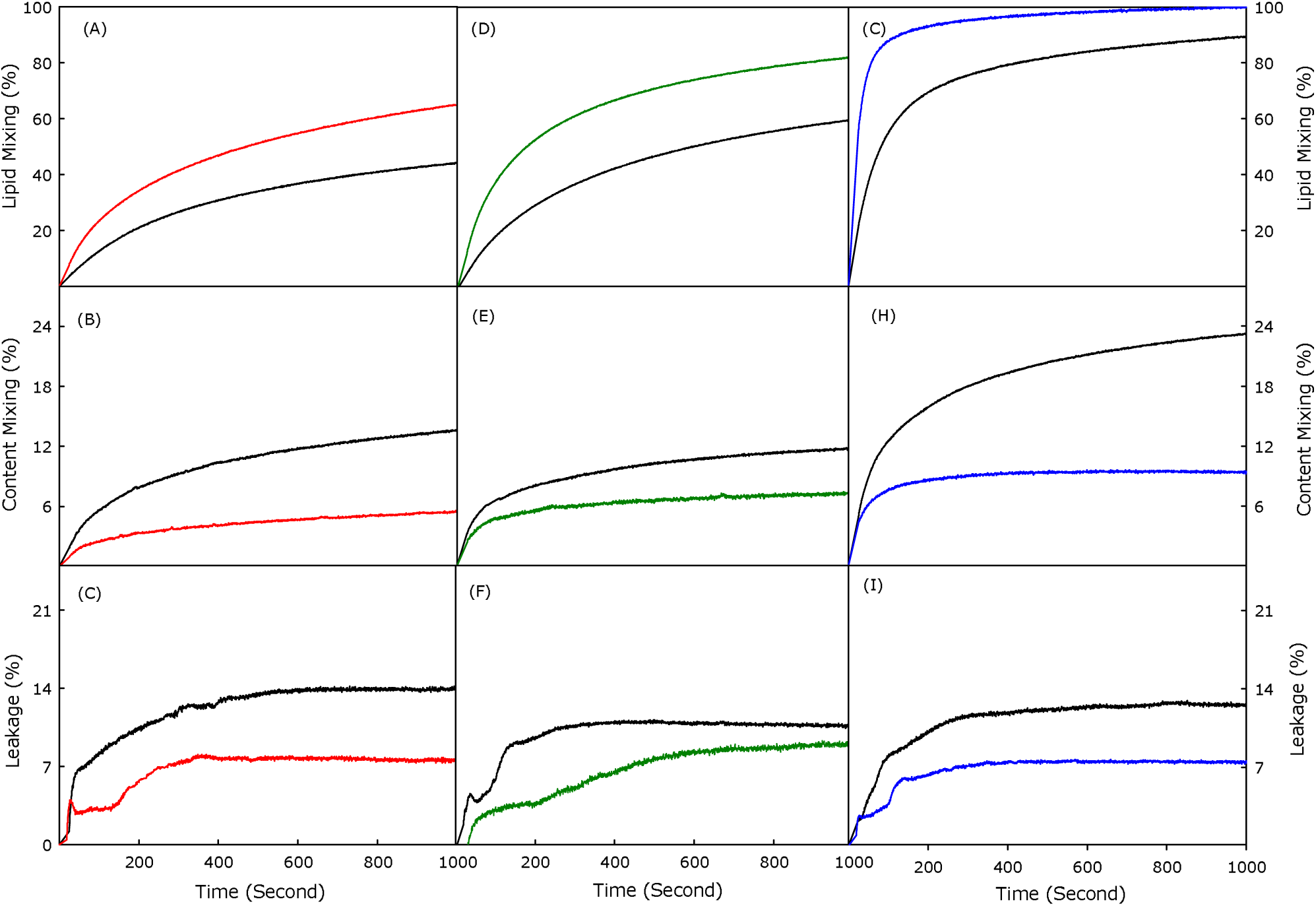
Plot of the effect of mGG-21 peptide on (A) lipid mixing, (B) content mixing, and C) content leakage in DOPC/DOPE/DOPG (60/30/10 mol%), (red) and (D) lipid mixing, (E) content mixing, and (F) content leakage in DOPC/DOPE/DOPG/CH (50/30/10/10 mol%) (blue) and (G) lipid mixing, (H) content mixing and (I) content leakage in DOPC/DOPE/DOPG/CH (40/30/10/20 mol%) (green) SUVs. Control (black) is the plot of all fusion observables in the absence of peptide. Fusion was induced by 6% (w/v) PEG at 37°C. Results are shown for a lipid-to-peptide ratio of 100:1. Measurements were carried out in 10 mM TES, 100 mM NaCl, 1 mM CaCl_2_, and 1 mM EDTA, pH 7.4 at a total lipid concentration of 200 µM. Data points shown are mean ± S.E of at least three independent measurements.

Cholesterol is an important constituent of the viral envelope.^51^ Therefore, we have performed similar experiments in membranes with varying cholesterol concentrations. Our results show that the mGG-21 peptide promotes LM by 25% and 7% in DOPC/DOPE/DOPG/CH (50/30/10/10 mol%) and DOPC/DOPE/DOPG/CH (40/30/10/20 mol%) membranes, respectively. However, the peptide inhibits CM by 42% and 62% in membranes containing 10 and 20 mol% of cholesterol. The kinetics of LM, CM, and L are shown in **Fig. 3**(**D-F**) and **Fig. 3**(**G-I**) for DOPC/DOPE/DOPG/CH (50/30/10/10 mol%) and DOPC/DOPE/DOPG/CH (40/30/10/20 mol%) membranes, respectively. The peptide inhibits pore formation in a similar fashion in cholesterol-containing membranes by stabilizing the hemifusion intermediates. It is important to mention here that the inhibitory efficiency of mGG-21 is much higher than that of GG-21 in all three membranes that we have studied.^42^ The inhibitory ability of GG-21 was around 20%. This signifies the importance of the addition of one extra WD repeat in the peptide sequence. The summary of LM, CM, and L results are shown in **Table 1**.

**Table 1.** The extent of lipid mixing, content mixing, and content leakage in the absence and presence of mGG-21 peptide in different lipid compositions. Data shown in the table are mean ± S.E of at least three independent measurements.

| Lipid Composition | Peptide | Lipid Mixing (%) | Content Mixing (%) | % of CM inhibition (Fusion Inhibition) | Content Leakage (%) |
| --- | --- | --- | --- | --- | --- |
| PC/PE/PG<br>(60/30/10 mol%) | Control | 59.8±2.4 | 15.2±0.5 | 59 | 10.4 ± 0.4 |
|  | mGG-21 | 75.5±3.0 | 6.2±0.2 |  | 7.3 ± 0.3 |
| PC/PE/PG/CH<br>(50/30/10/10 mol%) | Control | 69.5±2.8 | 12.3±0.4 | 42 | 11.1± 0.4 |
|  | mGG-21 | 87.2±2.9 | 7.1±0.2 |  | 9.4 ± 0.3 |
| PC/PE/PG/CH<br>(40/30/10/20 mol%) | Control | 93.0±3.8 | 24.2±0.9 | 62 | 12.4 ± 0.5 |
|  | mGG-21 | 100±4.0 | 9.3±0.3 |  | 7.5 ± 0.5 |

### Fusion Inhibitory effect of GG-21 versus mGG-21 peptide

To compare the fusion inhibitory ability of mGG-21 with its parent peptide GG-21 we have plotted the extent of inhibition in three different membranes with varying lipid compositions in **Fig. 4**. Our results demonstrate that the ability of mGG-21 to inhibit fusion is significantly higher than its native counterpart, which justifies the importance of WD repeat in designing better fusion inhibitors. Moreover, the WD repeat was located at the center of the GG-21 sequence whereas in mGG-21 the WDs are located close to two terminals of the peptide. This underscores the importance of WD at the terminal position of the inhibitory peptide. We have previously demonstrated that peptides with terminal WD exhibit superior inhibitory capacity in comparison to those with WD in the middle of the sequence.^40, 43^ It is well known that tryptophan and aspartic acid are located at the interface of the membrane because of their mixed polarity. Therefore, it is assumed that the WD repeat would act as an anchor to the membrane interface, and the rest of the sequence can be partitioned at the acyl chain region of the membrane. The anchoring is essential to wrap the membrane with the peptide. Coronin 1 is also known as the tryptophan-aspartic acid coat protein because it has multiple WD repeats that aid in coating the phagosome.^45^

**Figure 4.**
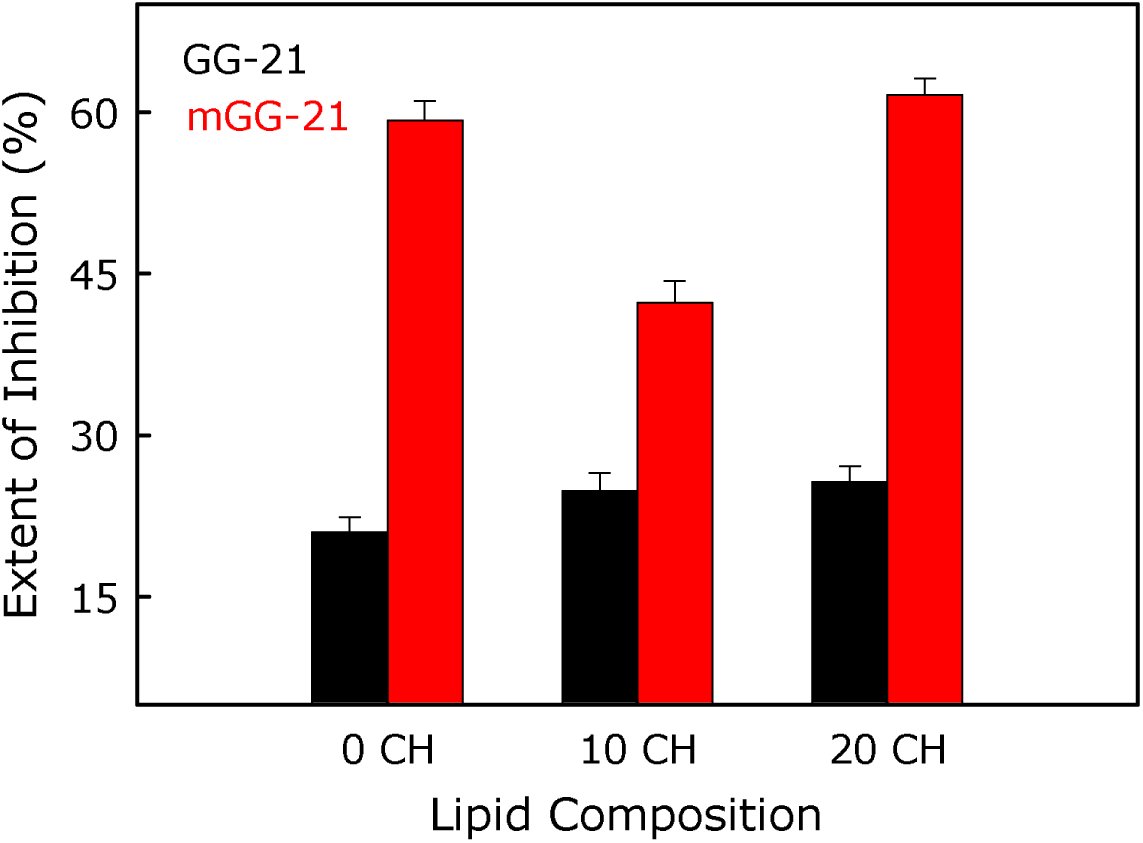
Comparison of percentage of fusion inhibition in terms of content mixing of GG-21 (black) and mGG-21 (red) peptides in DOPC/DOPE/DOPG (60/30/10 mol%), (0 CH) DOPC/DOPE/DOPG/CH (50/30/10/10 mol%) (10 CH) and DOPC/DOPE/DOPG/CH (40/30/10/20 mol %) (20 CH) SUVs. Data points shown are mean ± S.E of at least three independent measurements.

### Effect of the mGG-21 peptide on the bilayer properties

We have exploited the fluorescence properties of diphenyl hexatriene (DPH) and its trimethylammonium derivative (TMA-DPH) to evaluate the effect of mGG-21 on the depth-dependent membrane organization and dynamics. TMA-DPH is located at the interfacial region due to its polar trimethylammonium group with an average distance of 10.9 Å from the center of the bilayer.^52^ On the contrary, hydrophobic DPH is located at the acyl chain region of the bilayer with an average distance of around 7.8 Å from the bilayer center.^52^ The fluorescent properties such as anisotropy and the lifetime of TMA-DPH and DPH have been widely used to scrutinize the organization and dynamics of the membrane bilayer at two different depths.

### Effect of the mGG-21 peptide on the interfacial and acyl chain regions of the membrane

We have measured the peptide-induced change in fluorescence anisotropy of TMA-DPH in DOPC/DOPE/DOPG (60/30/10 mol%), DOPC/DOPE/DOPG/CH (50/30/10/10 mol %), and DOPC/DOPE/DOPG/CH (40/30/10/20 mol%) membranes in the presence of mGG-21 peptide. The lipid-to-peptide ratio was maintained at 100:1 as the fusion experiments were carried out at this lipid-to-peptide ratio. The fluorescence anisotropy of the interfacial probe, TMA-DPH, in different membranes in the presence and absence of peptide mGG-21 have been evaluated, and results are shown in **Fig. 5A**. From our results, it is evident that the fluorescence anisotropy of TMA-DPH in the presence of mGG-21 decreases in all three membranes. The decrease in fluorescence anisotropy of TMA-DPH suggests that the peptide disorders or fluidizes the interfacial region irrespective of the cholesterol content of the membrane. However, the interfacial ordering increases with increasing cholesterol content in the absence of mGG-21. This could be attributed to the inherent property of cholesterol to rigidify membranes.^53, 54^ We have further evaluated the interfacial polarity of the membrane in the absence and presence of mGG-21 by measuring the fluorescence lifetime of TMA-DPH in DOPC/DOPE/DOPG/CH ((100-x)/30/10/x mol %) membranes and the results are shown **Fig. 5B**. It is observed that the fluorescence lifetime of TMA-DPH partially increases in the presence of peptide in all lipid compositions indicating the partial reduction of interfacial polarity in the presence of the peptide. This reduction of polarity could be due to the exclusion of some water molecules by a non-polar peptide at the membrane interface.^39, 40^ We have calculated the apparent rotational correlation time of TMA-DPH from the fluorescence anisotropy and lifetime data using the Perrin equation^55^ (equation 7 in Supporting Information) to understand the effect of the peptide on membrane ordering more precisely. The results of the apparent rotational correlation time of TMA-DPH in different membranes in the presence and absence of mGG-21 have been shown in **Fig. 5C**. The apparent rotational correlation time data support the fluorescence anisotropy results. Overall, both the fluorescence anisotropy and apparent rotational correlation time results demonstrate that mGG-21 disorders the interfacial region. This peptide-induced disordering is correlated to the enhancement of lipid mixing in our fusion assay. The activation energy required to form the hemifusion state reduces in the membrane where the interfacial region is loosely packed. To evaluate the effect of the peptide on the order and polarity of the acyl chain region we have measured the fluorescence anisotropy and lifetime of DPH, a probe that partitions at 7.8 Å from the center of the bilayer. It is evident from the fluorescence anisotropy data (**Fig. 5D**) that the peptide orders the acyl chain region of the membrane at all lipid compositions. The ordering at the acyl chain region is often correlated to the inhibition in pore opening.^41, 42, 56^ The mGG-21-induced ordering supports the inhibition of content mixing, i.e., pore opening in membranes containing different amounts of cholesterol. Nevertheless, the fluorescence lifetime of DPH remains unchanged in the presence of mGG-21(**Fig. 5E**), suggesting that the peptide does not influence the acyl chain polarity. Generally, the peptide-induced enhancement of acyl chain polarity through water percolation is attributed to the enhancement of pore formation.^57^ For a better understanding of acyl chain ordering, we have calculated the apparent rotational correlation time using the Perrin equation in three membranes (**Fig. 5F**). The apparent rotational correlation time results support the fluorescence anisotropy values. Therefore, it can be hypothesized that the mGG-21 peptide blocks pore formation by ordering the acyl chain region of the membrane. We observed similar results for the GG-21 peptide.^42^ Consequently, our results suggest a strategy to design peptide-based fusion inhibitors with a property to order the acyl chain region.

**Figure 5.**
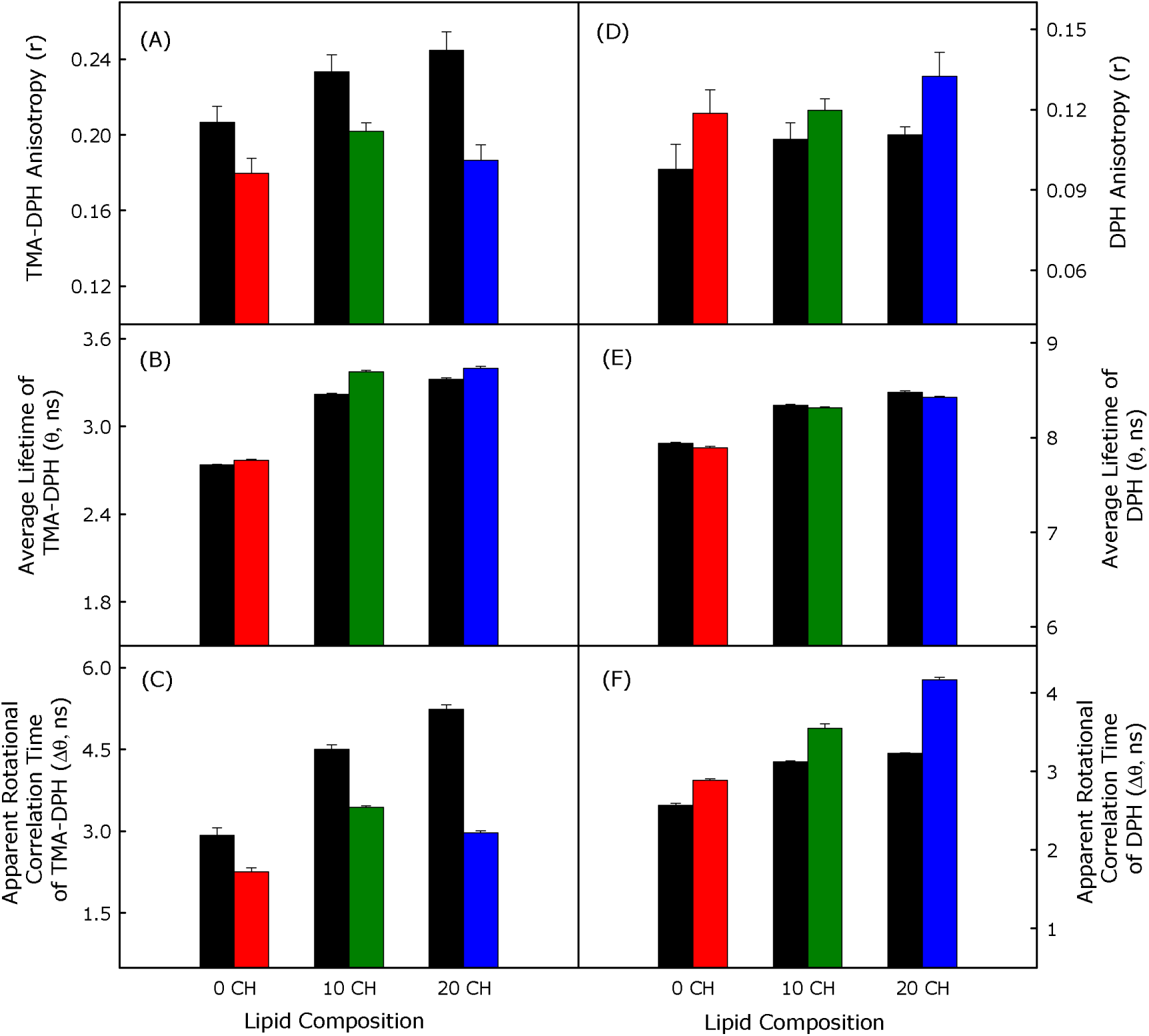
The plot of mGG-21 peptide-induced changes in the average (A) TMA-DPH anisotropy, (B) TMA-DPH fluorescence lifetime, (C) apparent rotational correlation time of TMA-DPH and (D) DPH anisotropy, (E) DPH fluorescence lifetime and (F) apparent rotational correlation time of DPH in DOPC/DOPE/DOPG (60/30/10 mol%) (red) DOPC/DOPE/DOPG/CH (50/30/10/10 mol %), (green) and DOPC/DOPE/DOPG/CH (40/30/10/20 mol%), (blue) SUVs in the presence and absence (black) of mGG-21. The lipid-to-peptide ratio was maintained at 100:1, at which we performed the fusion experiments. All the measurements were carried out in 10 mM TES, 100 mM NaCl, 1 mM CaCl_2_, and 1 mM EDTA, pH 7.4 at 37 °C, and a total lipid concentration of 200µM. Data points shown are mean ± S.E. of at least three independent measurements.

### Antiviral effect of the mGG-21

Since fusion plays a crucial role in the infection cycle of enveloped viruses and mGG-21 inhibits membrane fusion, we wanted to evaluate whether mGG-21 can inhibit viral infection, presumably by inhibiting endosome-virus fusion. To test this, we evaluated the antiviral activity of mGG-21 against H3N2 and H1N1 strains of the influenza virus. We infected A549 cells with X-31 (H3N2) and WSN (H1N1) strains of influenza virus in the absence and presence of varying concentrations of mGG-21 peptide. A549 cells are routinely used as a preclinical model for influenza infection. **Figure 6** shows that the influenza virus-induced severe cytopathic effects (CPE) after 48 hours of infection are significantly reduced in the presence of 50 µM mGG-21 peptide.

**Figure 6.**
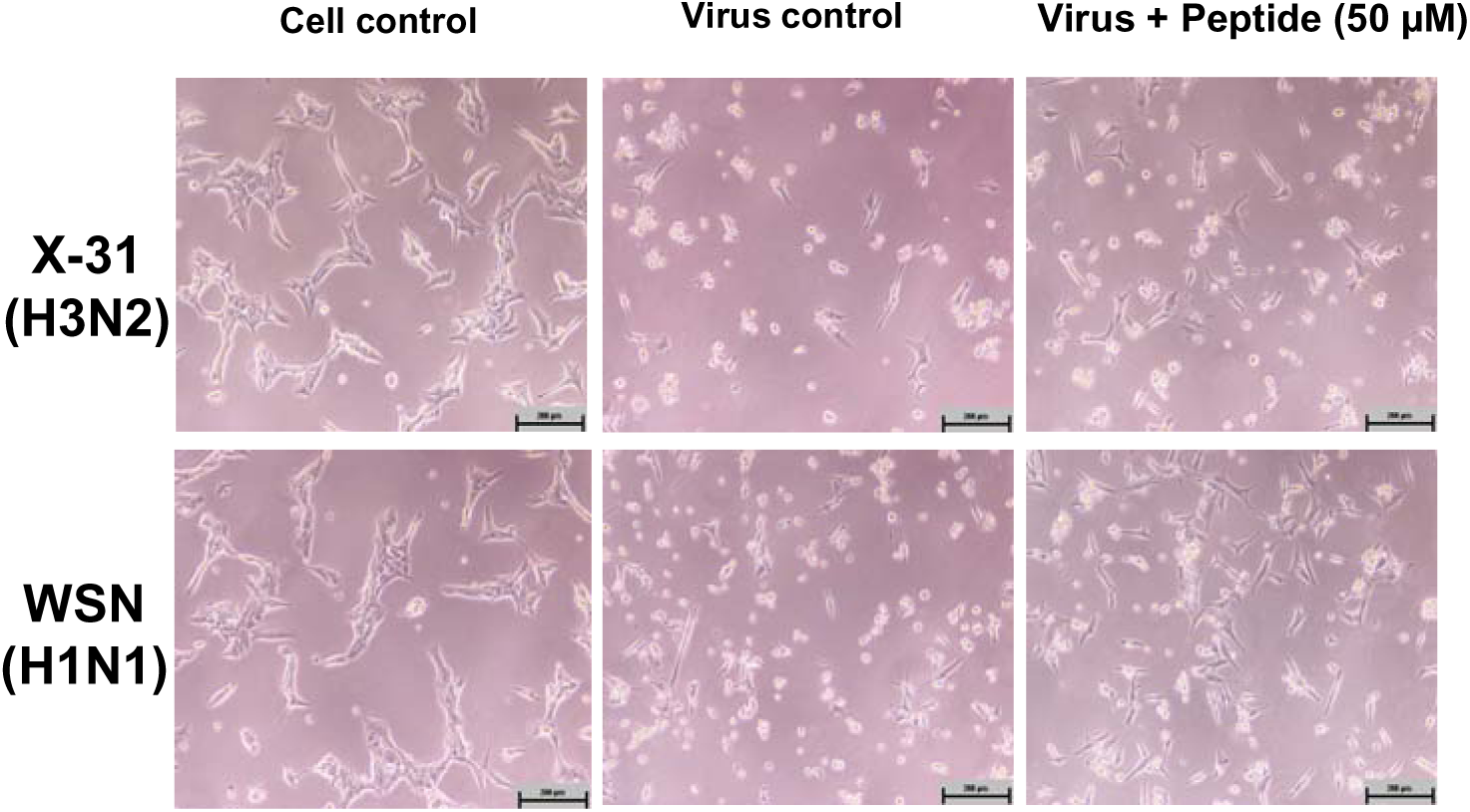
*In vitro* inhibitory activity of mGG-21 against H3N2 /X-31 (upper panel) and H1N1 /WSN (lower panel) strains of influenza virus infection. Light microscopy observation of a reduction in virus-induced cytopathic effect in the presence of mGG-21 in A549 cells after 48 hours of infection. A549 cells were infected with X31 (H3N2) and WSN (H1N1) influenza viruses in the absence and presence of 50 µM of mGG-21 peptide. Scale bar 200 µm. See the Materials and Methods section for details.

To quantify the inhibitory effect of mGG-21, we evaluated the post-infection cell viability by MTT assay in the presence of varying amounts of mGG-21. **Figure 7** shows a dose-dependent reduction in infection (i.e., increase in cell viability) of X-31 (H3N2) and WSN (H1N1). We quantified the post-infection cell viability as a normalized reduction in cytopathic effect (see Methods section for details and reference 43). **Figure 7 (C) and (D)** show dose-response curves of inhibition of X-31 (H3N2) and WSN (H1N1) by mGG-21 in cells. Although the reduction in cytopathic effect is comparable, the EC_50_ of X31 inhibition is ∼ 2.58 µM, and the EC_50_ of WSN inhibition is ∼29.39 µM. It is important to mention that the mGG-21 did not show any toxicity in A549 cells up to 50 µM (Supporting Information).

**Figure 7.**
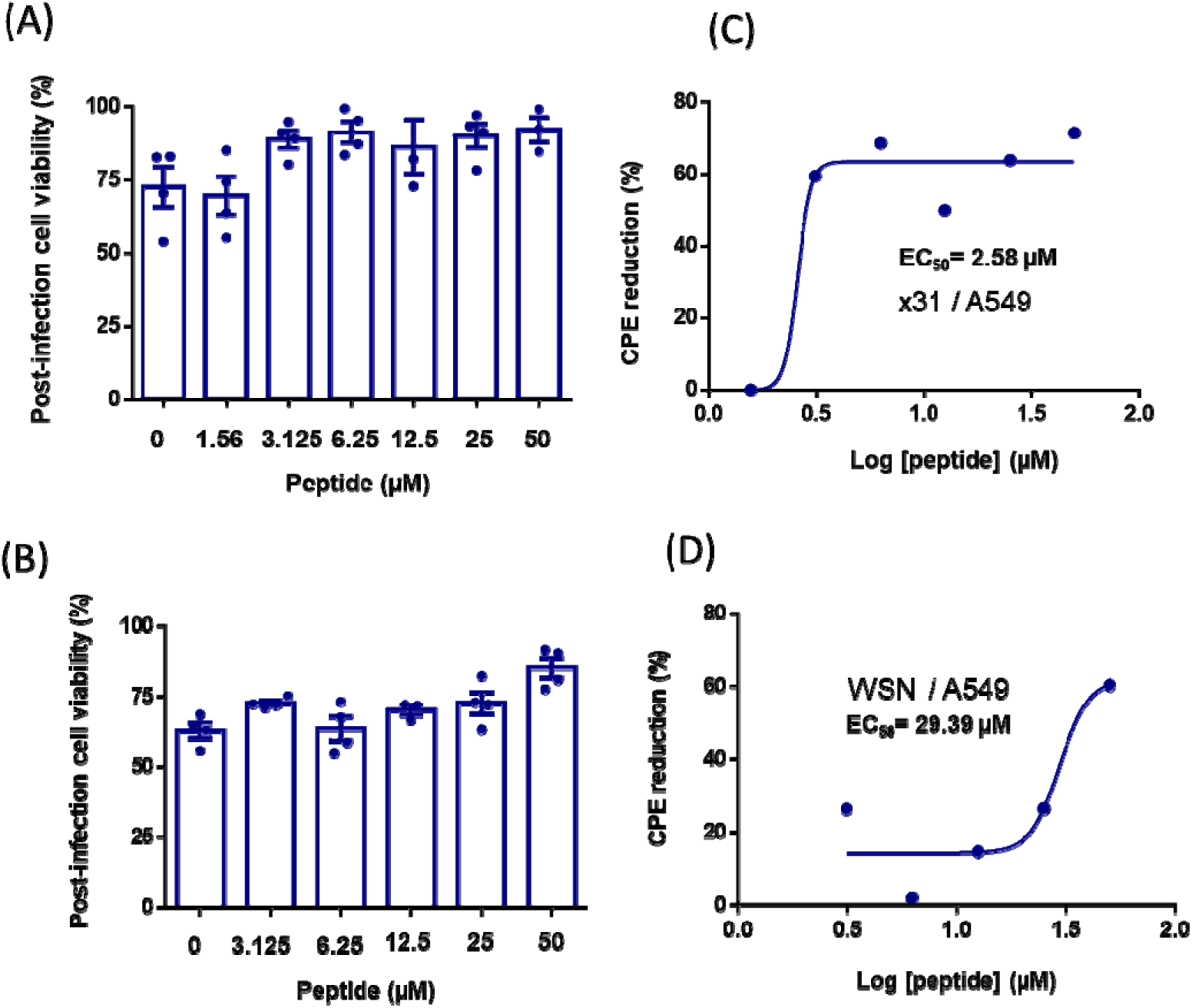
*In vitro* anti-influenza activity of mGG-21 against X31 and WSN. The effect of mGG-21 on the post-infection cell viability of A549 cells infected with (A) X31 and (B) WSN after 48 hours of infection was determined by the MTT assay. Cell viability was normalized to the control and represents the mean ± SE of at least 3 sets. Protection from CPE was normalized as described in the methods section and fitted to a dose-response curve for (C) x-31 and (D) WSN. See the method section for details.

To validate the reduction in infection of Influenza A Virus (IAV) by mGG-21, we carried out immunofluorescence staining with an anti-M1 (an Influenza viral protein) antibody of WSN-infected A549 cells in the presence and absence of mGG-21. A significant reduction of infected cells in the presence of 50 µM mGG-21 was observed (**Figure 8**). Next, we estimated the expression level of M1 in the absence and presence of mGG-21 peptide by Western blotting. **Figure 9** shows a reduction of M1 level compared to the virus control in A549 cells when infected by X-31 and WSN. Taken together, our findings demonstrate that the mGG-21 peptide inhibits X31 and WSN influenza virus infection in A549 cells.

**Figure 8.**
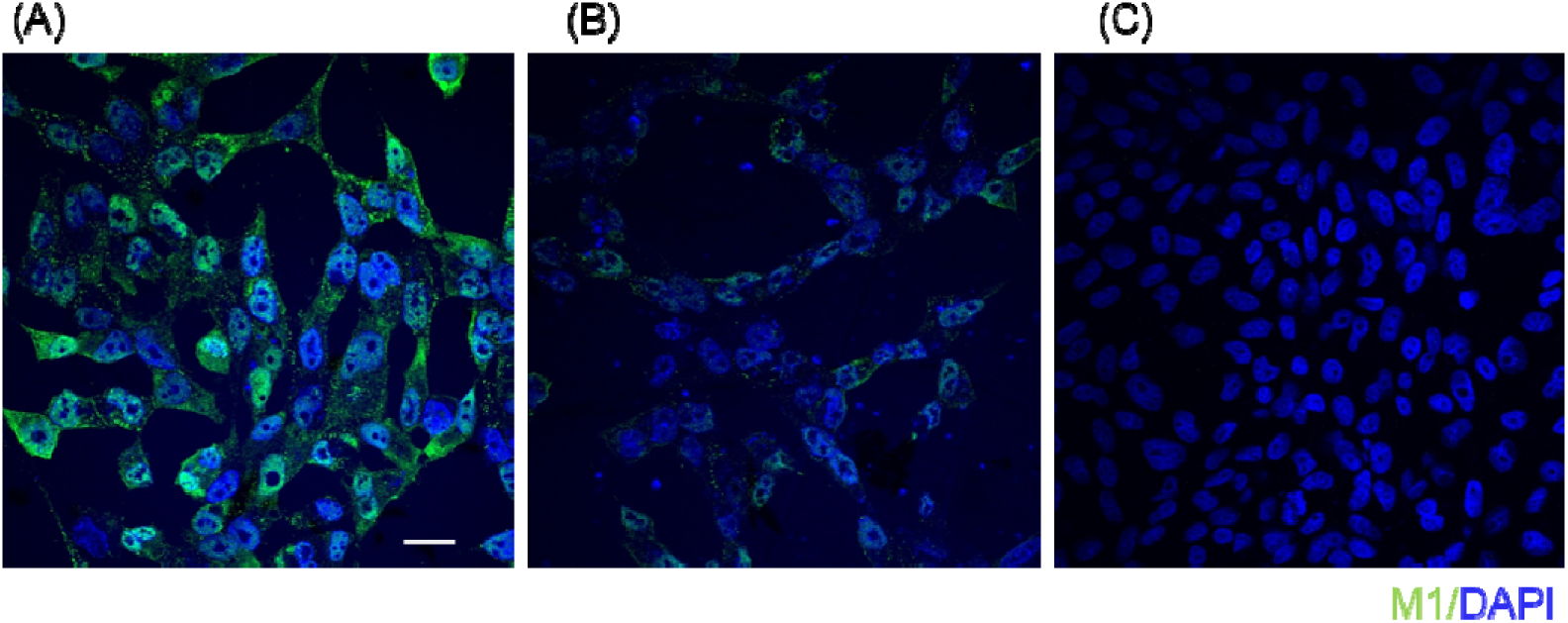
Imm unostaining of M1 in WSN-infected A549 cells. (A) Virus control (in the absence of mGG-21), (B) in the presence of mGG-21. (C) Cell control. A549 cells were infected with WSN (H1N1) influenza viruses in the absence and presence of 50 µM of mGG-21 peptide, fixed after 15 hours, and stained with a polyclonal anti-M1 antibody. Cells were counterstained with DAPI. Scale bar 20 µm. See the method section for details.

**Figure 9.**
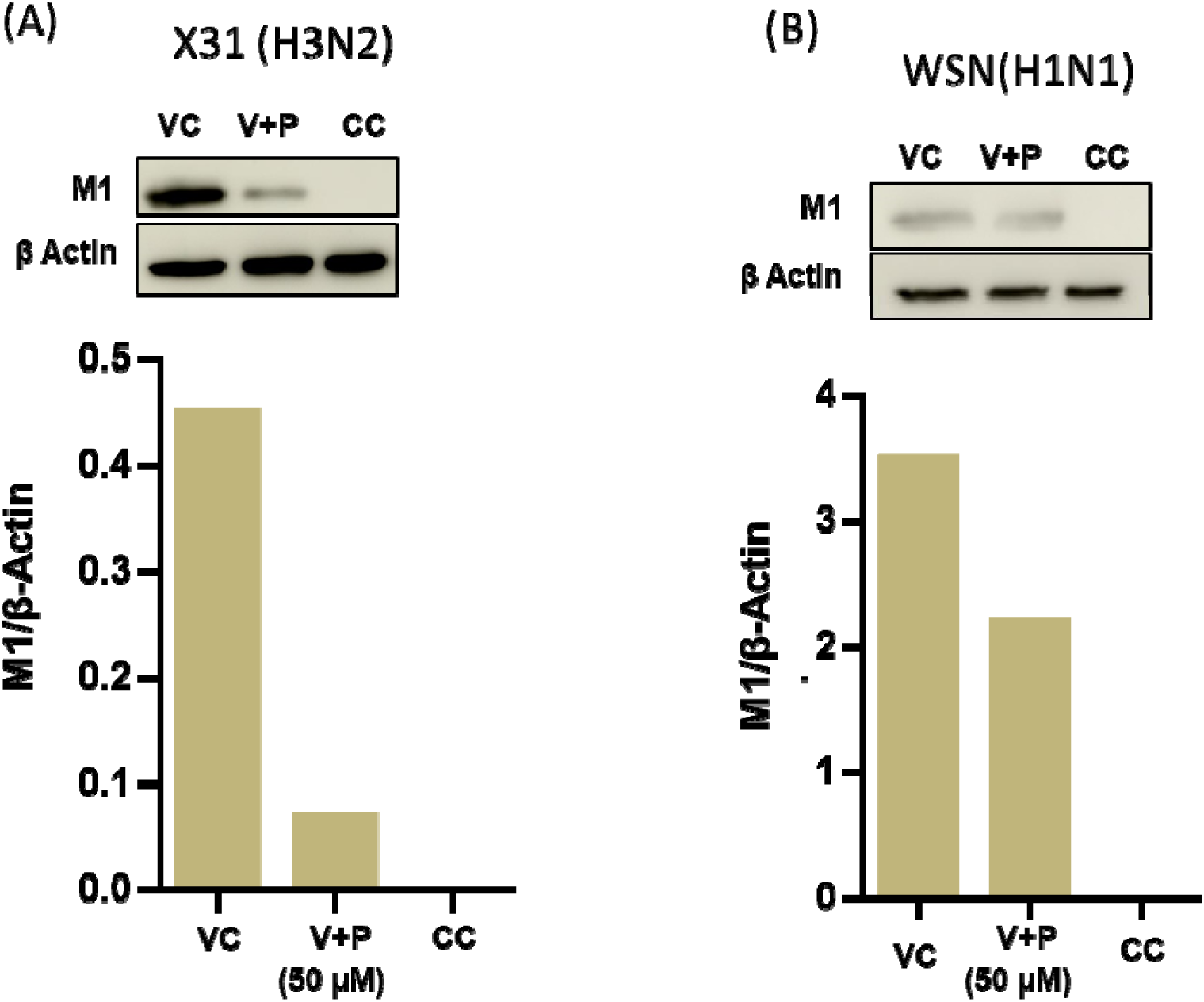
Reduction in viral protein (M1) expression. M1 expression levels in A549 cells in the presence of 50 µM mGG-21; (A) infected with X-31 (H3N2) and (B) infected with WSN(H1N1). M1 expression level has been evaluated after 48 hours of infection in the absence and presence of the mGG-21 peptide using Western Blot. β-actin was taken as a loading control.

If the antiviral effect of mGG-21 is due to its ability to inhibit virus-endosome fusion without specifically targeting a particular protein in influenza, it may also inhibit the fusion of other viruses and prevent subsequent infections. To explore this hypothesis, we have examined the antiviral effects of mGG-21 against chikungunya virus (CHIKV). Chikungunya virus transfers its genome to the host cells by fusion of the viral envelope with endosomal membranes, akin to Influenza, however, with a class II fusion protein, unlike Influenza Virus, which has a Class I fusion protein.^20^ Our results show that mGG-21 displays a moderate but consistent antiviral effect against CHIKV infection in BHK-21 cells. The mGG-21 showed a 50% inhibition in virus infection at a 50 µM concentration consistently in repeated experiments (**Figure 10**). Combining the antiviral data with the results observed from model membrane studies, it is likely that mGG-21 inhibits influenza and chikungunya infection by preventing virus-endosome fusion. Nevertheless, our findings do not rule out the possibility of other mechanisms for the antiviral action of mGG-21.

**Figure 10.**
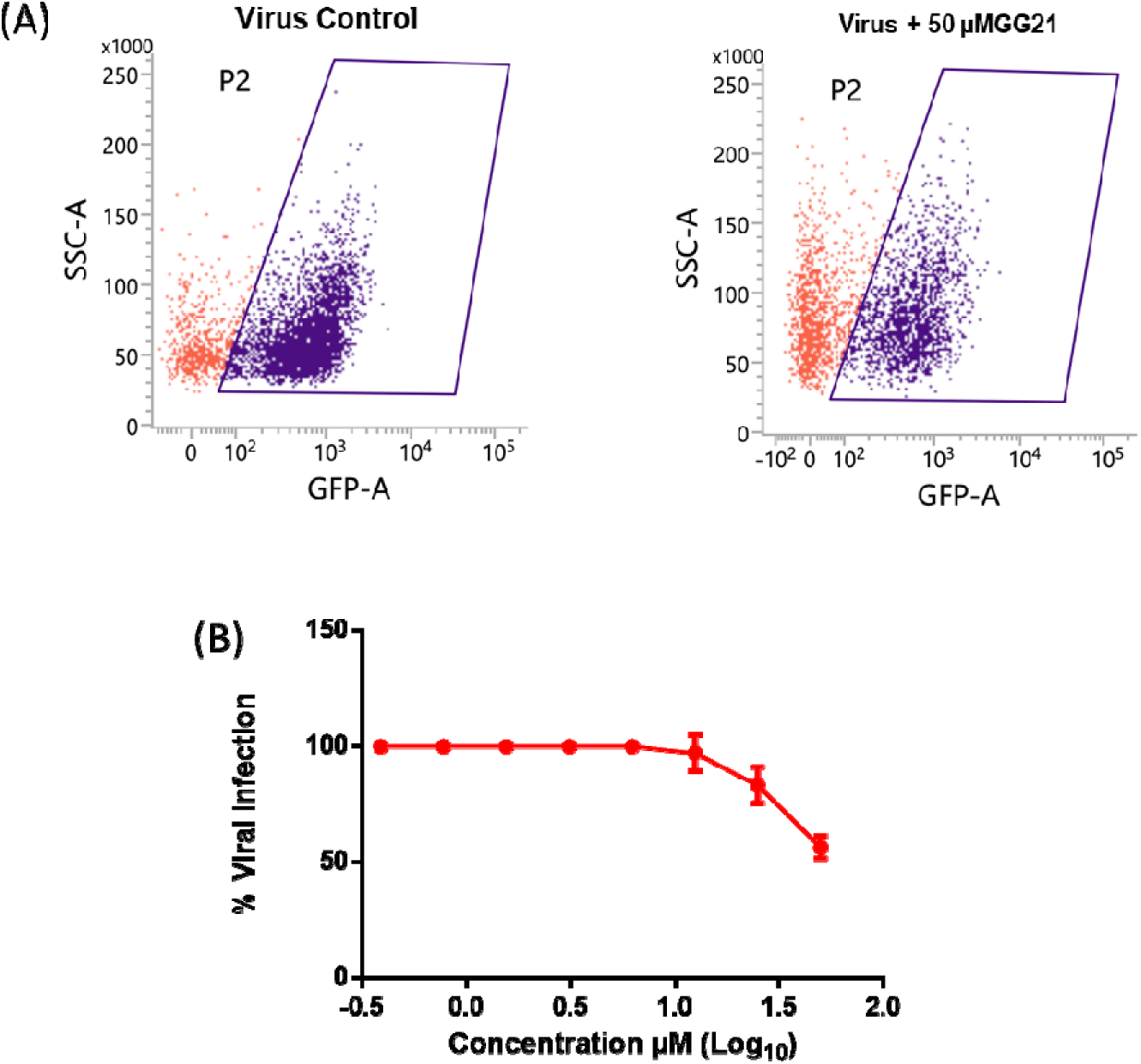
Effect of mGG-21 on Chikungunya virus infection in BHK-21 cells. (A) Representative flow cell scatter diagrams show the reduction in CHIKV-GFP-infected Vero cells in the presence of 50 µM mGG-21 and (B) the dose-dependent reduction in Chikungunya infection by mGG-21.

## Discussion

Due to antigenic shift and antigenic drift, pathogenic viruses escape the immune surveillance and pose a looming threat of future pandemics. Therefore, there is a clear need to develop broad-spectrum strategies beyond the ‘one-bug-one-drug’ paradigm to combat growing numbers of diverse emerging and re-emerging viral pathogens.^38^ Targeting the viral fusion pathway may lead to a novel broad-spectrum antiviral strategy. It is already known that Interferon-induced transmembrane protein 3 (IFITM3) exerts its broad-range antiviral activity by inhibiting virus-endosome fusion.^58^ In this paper, we posit that viral fusion and hence infection can be inhibited by appropriately designed peptides that target the participating viral (lipid) envelope and host cellular membranes (plasma membrane or endosomal membranes) without explicitly targeting a particular viral fusion protein. To develop such broad-spectrum fusion inhibitors, we have utilized the successful defence mechanism of the mycobacterium species. Phagosomes containing living mycobacteria have been shown to avoid lysosomal fusion by recruiting and retaining a phagosomal protein, coronin 1, which contains conserved Trp-Asp (WD) repeats on the phagosomal surface. We have designed a novel peptide, mGG-21, that inhibits protein-independent model membrane fusion. The mGG21 modulates the biophysical properties of the participating membranes to impede fusion. Utilizing complementarity approaches, including Western blotting and FACS, we have demonstrated that mGG-21 inhibits Influenza and Chikungunya virus infection in cells. Although they infect cells via the endosomal fusion pathway, the influenza virus belongs to the *Orthomyxoviridae* family with class I fusion protein, chikungunya belongs to the *Togaviridae* family containing class II fusion protein.^59^ Since mGG21 inhibits two different classes of enveloped virus infection, it must be through a common mechanism, likely through inhibition of virus-endosome fusion. However, our results do not rule out alternative mechanisms such as inhibition of endocytosis.

Our results rationalize the importance of lipid-peptide interplay in modulating membrane organization and dynamics in designing WD repeat-containing peptide-based fusion inhibitors. The peptide-induced ordering of the acyl chain region could be successfully exploited to develop peptide-based fusion inhibitors. The WD-repeat containing mutated peptide from coronin 1 successfully inhibits the pore opening in the model membrane fusion assay by stabilizing the hemifusion intermediate. The inhibitory efficacy of mGG-21 is significantly higher than the other peptide-based fusion inhibitors developed utilizing coronin 1 as a template.^40–43^ The depth-dependent membrane ordering results show that the peptide orders the acyl chain region and does not allow water percolation in this region, thereby stabilizing the hemi-fusion stalk (or diaphragm). This leads to inhibition of fusion pore formation. In our experiment, enveloped viruses like influenza and chikungunya are introduced to cells in the presence of mGG21. The peptides might interact with viral envelopes in a manner that impedes fusion with the endosomes and stabilizes the hemifusion structures, consequently inhibiting infection.

## Conclusions

Our results rationalize the importance of lipid-peptide interplay in modulating membrane organization and dynamics in designing WD repeat-containing peptide-based fusion inhibitors. The peptide-induced ordering of the acyl chain region could be successfully exploited to develop peptide-based fusion inhibitors. The WD-repeat containing mutated peptide from coronin 1 successfully inhibits the pore opening in the model membrane fusion assay by stabilizing the hemifusion intermediate. The inhibitory efficacy of mGG-21 is significantly higher than the other peptide-based fusion inhibitors developed utilizing coronin 1 as a template. The depth-dependent membrane ordering results show that the peptide orders the acyl chain region and does not allow water percolation in this region, resulting in the inhibition of pore formation. The peptide was further tested for its antiviral activity against Influenza (X-31 (H3N2) and WSN (H1N1) strains) and Chikungunya viruses. It was observed that infection of both influenza and Chikungunya virus was reduced significantly when the cells were incubated with 50 µM of mGG-21 peptide. Additionally, mGG-21 has hardly any cytotoxic effect at this concentration (**Figure S1**). Overall, our approach to designing peptide-based fusion inhibitors or entry inhibitors might be a helpful strategy against the pandemics caused by the emerging and remerging viral infections.

## Author contributions

S. P. carried out the spectroscopic and model membrane experiments, analyzed the data. B.Q., V,V., and S.G. performed the biochemical and virology experiments for Influenza and Chikungunya. S.P. and B.Q. wrote the initial draft of the manuscript. H.C., C.D.M., and S.H. conceptualized and supervised the work, acquired funding, and wrote and edited the final manuscript.

## Acknowledgments

This work was supported by the Core Research Grant (CRG/2021/001515) of the Science and Engineering Research Board (SERB), the SERB-Science and Technology Award (STR/2021/000029) for Research, New Delhi, to HC, Ramalingaswami Re-entry Fellowship (BT/RLF/Re-entry/11/2020) to S.H., and SERB-Start-up Research Grant to CM. H.C. thanks the University Grants Commission for the UGC-Assistant Professor position and, SERB for the fellowship. S.P. and B.Q. thank the University Grants Commission and CSIR for the Research Fellowships, respectively. S.H. and C.D.M. thank the Division of Virus Research and Therapeutics, CSIR-CDRI, for providing the infrastructure. X-31 Influenza A virus was a kind gift from Dr. Joshua Zimmerberg from NIH, Bethesda. WSN (H1N1) virus was a kind gift from Dr. Arindam Mandal, IIT Kharagpur. H.C. thanks DST, New Delhi, and UGC for providing the instrument facility to the School of Chemistry, Sambalpur University, under the FIST and DRS programs, respectively. We thank the Sophisticated Analytical Instrument Facility (SAIF), CSIR-CDRI, for providing us with the Confocal Microscopy (Intravital facility) and Flow Cytometry facilities. We thank members of the Chakraborty and Haldar laboratories for their comments and discussions.

## TOC Graphic

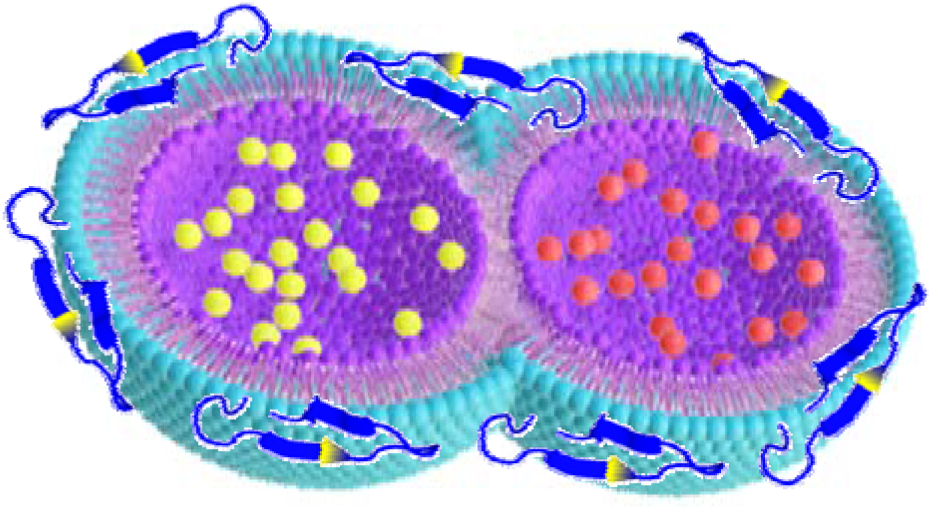

## Supporting Information

### MATERIALS AND METHODS

#### Materials

1,2-dioleoyl-sn-glycero-3-phosphocholine (DOPC), 1,2-dioleoyl-sn-glycero-3-phosphoethanolamine (DOPE), cholesterol (CH), 1,6-Diphenyl-1,3,5-hexatriene (DPH) and trimethyl derivative of 1,6-Diphenyl-1,3,5-hexatriene (TMA-DPH) were procured from Avanti Polar Lipids (Alabaster, AL). N-(7-nitrobenz-2-oxa-1,3-diazol-4-yl)-1,2-dihexadecanoyl-sn-glycero-3-phosphoethanolamine (NBD-PE), and Lissamine rhodamine B 1,2-dihexadecanoyl-sn-glycero-3-phosphoethanolamine (Rh-PE) were purchased from Thermo Fisher Scientific (Waltham, MA). Triton X-100 (TX-100) and poly (ethylene glycol) of molecular weight 7000-9000 gm (PEG 8000) were procured from Sigma Aldrich (St. Louis, MO). N-[Tris(hydroxymethyl)-methyl]2-2-aminoethane sulfonic acid (TES) was obtained from Alfa Aesar (Haverhill, MA). Calcium chloride and ethylenediaminetetraacetic acid (EDTA) were obtained from Merck (India), whereas sodium chloride was obtained from Fisher Scientific (India). UV-grade solvents such as chloroform and methanol have been procured from Spectrochem (India). All the chemicals used in the work are of 99% purity. Water was purified through an Adrona Crystal (Latvia) water purification system. All chemicals utilized in this work have more than 99% purity.

#### Peptide Synthesis

The mGG-21 peptide sequence is GC**WD**VI**L**V**AV**VGTGAA**VWD**LG, which was chemically synthesized and purified by Biochain Incorporated (New Delhi, India). The peptides were synthesized by the solid phase approach using Fmoc chemistry,^1, 2^ which is discussed elsewhere. Mass spectrometry has been utilized to characterize the peptide that was isolated via HPLC. Small aliquots of the peptide stock solutions were added to the vesicle suspensions after they were prepared in DMSO. The DMSO concentration was always less than 1% (v/v), and it had no discernible impact on membrane shape or fusion.^3^

#### Preparation of Vesicles

Using the sonication method, small unilamellar vesicles (SUVs) were prepared from either DOPC/DOPE/DOPG (60/30/10 mol %), DOPC/DOPE/DOPG/CH (50/30/10/10 mol %), and DOPC/DOPE/DOPC/CH (40/30/10/20 mol %). Lipids at this appropriate molar ratio, in chloroform, were dried overnight in a vacuum desiccator. For uniform dispersion of lipids, the dried films were hydrated and vortexed in the assay buffer for 1 h. The column (Sephadex G-75) and experimental buffer were comprised of 10 mM TES, 100 mM NaCl, 1 mM EDTA, and 1 mM CaCl_2_ at pH 7.4. Then, to prepare SUVs, the hydrated lipids were sonicated using a Dr. Hielscher model UP100H (Germany) probe sonicator, as documented previously.^4, 5^ Sonicated samples were centrifuged at 15,300 g for 10 minutes to remove aggregated lipids and titanium particles. All lipid mixing, content mixing, and leakage experiments were performed with 200 μM lipid. Using dynamic light scattering (Litesizer DLS 500, Anton Paar), the average hydrodynamic radii of the vesicles were determined to be 50-60 nm with a polydispersity index of less than 0.2 (data not shown). Small aliquots of the probe were added from their respective stock solutions prepared in DMSO into working solutions. The DMSO content was always less than 1% (v/v), and this small quantity of DMSO was established not to produce any detectable effect on membrane structure.

#### Steady-State Fluorescence Anisotropy Measurements

Steady-state fluorescence anisotropy measurements were performed using a Hitachi F-7000 (Japan) spectrofluorometer with a 1cm path-length quartz cuvette. TMA-DPH and DPH were excited at 360 nm, and their emissions were monitored at 428 nm. In all measurements excitation and emission slits with a nominal bandpass of 5 nm were used. All measurements were conducted in a buffer containing 10 mM TES, 100 mM NaCl, 1 mM CaCl_2_, and 1 mM EDTA, pH 7.4 at 37 ^°^C. Anisotropy values were calculated using the following equation^6^

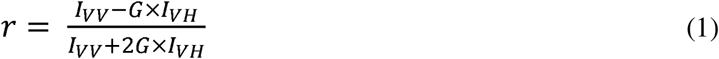

where, G = I_HV_/I_HH_, (grating correction or G-factor), I_VV_ and I_VH_ are the measured fluorescence intensities with the excitation polarizer vertically oriented and the emission polarizer vertically and horizontally oriented, respectively.

#### Lipid Mixing Assay

The transfer of lipid (Lipid Mixing) during PEG-mediated vesicle fusion was examined based on the change in FRET efficiency between FRET lipid pairs: NBD-PE (donor) and Rh-PE (acceptor).^5, 7^ FRET dilution as a function of time is reflected as a marker to measure the kinetics of lipid transfer (mixing) between two vesicles. We prepared a set of vesicles comprising FRET lipid pairs in equal concentration (0.8 mol %) and indicating maximum FRET. The probe-containing vesicles were mixed with probe-free vesicles at a ratio of 1:9.^8^ 6% (w/w) PEG was added to 200 μM lipid vesicles to initiate the lipid mixing process, which was measured by observing the reduction in FRET efficiency via a rise in donor intensity. To evaluate the effect of inhibitor peptide, 2 μM peptide was added to the mixture of probe-containing and probe-free (1:9) vesicles 10 mins before the addition of PEG to monitor the lipid mixing. The emission intensity of the donor (NBD-PE) was examined with Hitachi F-7000 (Japan) spectrofluorometer, keeping the excitation and emission wavelengths fixed at 460 nm and 530 nm, respectively. Throughout the experiment, a minimum slit of 5 nm was used on both the excitation and emission sides. Each experiment was repeated at least three times. The percentage of lipid mixing was calculated using the following equation^7^

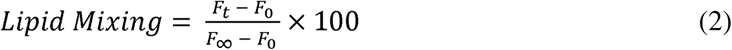

where ‘F_0_’, ‘F_t_’, and F_∞_’ are the fluorescence intensities at the zeroth time, time = t and time = ∞, respectively. F∞ has been measured in the presence of TX-100, which is considered the complete mixing of lipids.

#### Content Mixing Assay

We have monitored the content mixing as proposed by Wilschut et al. using Tb^3+^ and DPA.^9, 10^ Vesicles were prepared either in 80 mM DPA or 8 mM TbCl_3_. The untrapped DPA and TbCl_3_ were removed from the external buffer of the vesicles using a Sephadex G-75 column equilibrated with assay buffer (10 mM TES, 100 mM NaCl, 1 mM EDTA,1 mM CaCl_2_ at pH 7.4). 6% (w/w) PEG was added to 200 μM lipid vesicles to induce content mixing. 2 μM peptide was added to a 200 μM mixture (1:1) of Tb^3+^ and DPA-containing vesicles 10 mins before the addition of PEG to monitor the content mixing. The content mixing was measured in terms of an increase in fluorescence intensity due to the formation of the Tb/DPA complex with time. The excitation and the emission wavelength for the content mixing assay were used as 278 nm and 490 nm, respectively. A minimum slit width of 5 nm in both the excitation and emission side was used for all measurements. Each experiment was repeated at least three times. The percentage of content mixing was calculated in the following way:

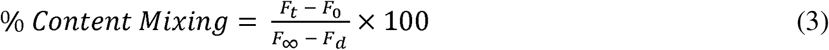

where ‘F_0_’ and ‘F_t_’ are the fluorescence intensities at zeroth time and time = t, respectively. ‘F_∞_’ and ‘F_d_’ have been calculated from the fluorescence intensity of the leakage sample at a zeroth time (which gives the highest intensity as all Tb are being complexed with DPA) and in the presence of detergent, respectively. ‘F_∞_’ implies the highest fluorescent signal when Tb^3+^ and DPA are in the same vesicular compartment and all Tb^3+^ are in a complex form with DPA. Upon detergent addition, Tb^3+^ and DPA will come out from the vesicular compartment and be diluted in the bulk solution, resulting in dissociation of the complex and minimum fluorescence intensity (F_d_).

#### Leakage Assay

The leakage assay was carried out by monitoring the decrease in fluorescence intensity of the vesicles containing both TbCl_3_ and DPA in the presence of PEG or PEG and peptide.^9, 10^ 8 mM TbCl_3_ prepared in 10 mM TES and 100 mM NaCl, pH 7.4, and 80 mM DPA which is prepared in 10 mM TES, pH 7.4, were co-encapsulated in the vesicles. The external TbCl_3_ and DPA were removed using a Sephadex G-75 column equilibrated with assay buffer (10 mM TES, 100 mM NaCl, 1 mM EDTA, 1 mM CaCl_2_ at pH 7.4).^8^ 6% (w/w) PEG was added to 200 μM lipid Tb^3+^ and DPA-containing vesicles to induce content leakage. To evaluate the effect of inhibitor peptide, 2 μM peptide was added to the vesicles 10 mins before the addition of PEG to monitor the content leakage. The maximum leakage (100%) was characterized by the fluorescence intensity of a co-encapsulated TbCl_3_/DPA vesicle treated with 0.1% (w/ v) Triton X-100. The excitation and the emission wavelength were fixed at 278 nm and 490 nm, respectively. Slits of 5 nm were used in both the excitation and emission sides throughout the experiment. Each experiment was repeated at least three times. The percentage of content leakage was calculated in the following way:

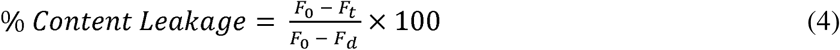

where ‘F_0_’ and ‘F_t_’ and ‘F_d_’ are the fluorescence intensities at the zeroth time, time = t, and in the presence of detergent, respectively.

#### Time-resolved Fluorescence Measurements

Fluorescence lifetimes were measured from time-resolved fluorescence intensity decays using the IBH 5000F Nano LED equipment (Horiba, Edison, NJ) with Data Station software in time-correlated single photon counting (TCSPC) mode, as mentioned earlier.^5, 10^ A pulsed light-emitting diode (LED) was applied as the excitation source. The LED generated an optical pulse at 340 nm (for exciting DPH and TMA-DPH), with a pulse duration of less than 1.0 ns, and was run at a rate of 1 MHz. The Instrument Response Function (IRF) was measured at the respective excitation wavelength using Ludox (colloidal silica) as a scatterer. Ten thousand photon counts were collected in the peak channel to improve the signal-to-noise ratio. All experiments were performed using emission slits with an 8-nm bandpass for DPH and TMA-DPH. Data were stored and analysed using DAS 6.2 software (Horiba, Edison, NJ). Fluorescence intensity decay curves were deconvoluted with the instrument response function and analysed as a sum of exponential terms as follows:

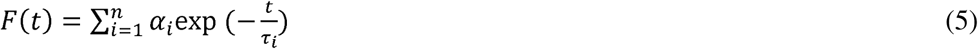

A considerable plot was found with a random deviation around zero with a maximum χ2 value of 1.2 or less. Intensity-averaged mean lifetimes τ_avg_ for n-exponential decays of fluorescence were calculated from the decay times and pre-exponential factors using the following equation^6^

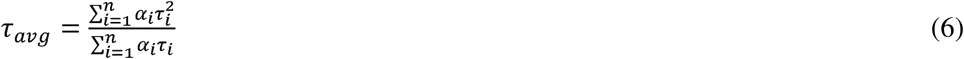

where α_i_ is the fraction that shows lifetime τ_i_.

#### Calculation of Apparent Rotational Correlation Time

Apparent rotational correlation time delivers information about the ease of rotational motion of fluorophores during their lifetime, which is affected by the rigidity of its immediate microenvironment. Therefore, we can acquire information about membrane viscosity in the interfacial as well as the hydrophobic region by monitoring the apparent rotational correlation time (θ_c_) of DPH and TMA-DPH from steady-state anisotropy and the average lifetime values using Perrin’s Equation^6^ as follows:

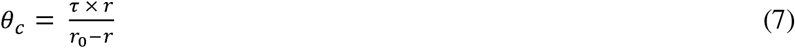

where ‘τ’ is the average lifetime of the probe and r is anisotropy. The constant r_0_ value was considered as 0.4 for DPH and TMA-DPH.^6^

#### Cell Line, Viruses, and Antibodies

A549 and BHK-21 cells were obtained from ATCC and were maintained in DMEM (GIBCO) with 10 % FBS (HIMEDIA) containing 100 μg/ml penicillin (HIMEDIA) and 100 μg/ml streptomycin (HIMEDIA). Cells were grown in a humidified atmosphere (95% humidity) at 37 °C and 5% CO_2_. Trypsin was obtained from HIMEDIA. The influenza A virus (strain X-31, A/Aichi/68, H_3_N_2_) and (strain WSN, H_1_N_1_) were kind gifts from Dr. Joshua Zimmerberg at NICHD/ NIH, Bethesda, and Dr. Arindam Mondal (IIT Kharagpur), respectively. A rabbit polyclonal anti-M1 antibody (GTX-127356) and a goat anti-rabbit IgG H&L [HRP] (ab6721). An HRP-conjugated anti-β actin (A3854) was obtained from Sigma. Green Fluorescent protein tagged chikungunya virus, CHIKV-GFP (181/25) was created using a reverse genetics platform (Dr. Chetan D Meshram, unpublished data).

#### Influenza Virus Infectivity and Peptide Efficacy Assay

X-31 (H_3_N_2_) and WSN (H1N1) Influenza viral infection was carried out in the A549 cell line. The *in vitro* antiviral effect of the peptide was evaluated by quantifying the reduction in the cytopathic effect induced by the Influenza A virus in A549 cells. 1.5 X 10^4^ (A549) cells were seeded in each well of a 96-well tissue culture plate and were allowed to grow as a monolayer for 24 hours. After adherence, the cells were incubated with different concentrations of the peptide for 48 hours at 37 °C and 5% CO_2_. After 48 hours, cell viability was assessed by MTT assay. For infection, the peptide (mGG21) was two-fold serially diluted (from a 10 mM stock in DMSO) in serum-free media (SFM) containing trypsin (2.5 µg/ml). Equal amounts of Influenza A virus from stock were pre-mixed with different amounts of the peptide in SFM. A549 cells were then infected with peptide-virus mixtures for 45 minutes at 37°C, 5% CO_2._ After that, the infection media was removed, followed by washing the cells with PBS to remove unbound virus, and then supplemented with growth media containing 0.1% FBS, 0.3% BSA, and trypsin (2.5 µg/ml) and incubated for 48 hours at 37°C, 5% CO_2_. After 48 hours, 10μl of MTT (SRL) (from a stock of 5 mg/ ml in PBS) was added to each well and the plate was incubated for 4 hours at 37°C, 5% CO_2._ After that, 100 μl of DMSO was added to each well to dissolve the formazan crystals produced in the process, and absorbance was measured at 570 nm. The wells containing only cells and cells infected with the virus were considered as cell control and virus control, respectively. Cell viability was determined by considering that the cell control absorbance corresponds to 100 % viability. The percentage of cytopathic effect (CPE) inhibition was defined as

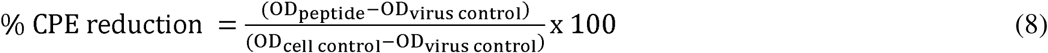

where OD_peptide_ is the absorbance in the presence of virus and peptide, OD_virus control_ is the absorbance in the presence of virus only and OD_cell control_ is the absorbance of cell control wells.^11^ Each O.D. value was background subtracted. Reduction in CPE is plotted as a function of peptide concentration and fitted to a dose-response model in GraphPad Prism 6 (San Diego, CA, USA), and 50% effective concentration (EC_50_) was extracted (see **Fig. 5**).

#### Western blot analysis

To determine the level of Influenza viral protein M1 expression, A549 cells were seeded in a 6-well plate at 0.25×10 ^6^ cells / well and infected with Influenza A virus in the absence and presence of varying amounts of mGG21 as described above. After 48 hours of infection, cells were observed for cytopathic effect (See Fig. 6 A). After removing the media, cells were washed with ice-cold PBS and scrapped in a lysis buffer (RIPA-HIMEDIA: 50 mM Tris-HCl pH 7.4, 1% Triton X-100, 1% sodium deoxycholate, 1 mM EDTA, 0.1% SDS) in the presence of a protease inhibitor cocktail (SIGMA). To prepare cell lysate, cells were subjected to sonication for 90 seconds (pulse 15, Amp1 60%) (Fisher Scientific) and then pelleted by centrifugation at 16000 RCF for 20 minutes at 4 °C. The supernatant was collected and stored at −20 °C. Total protein concentration was determined by a bicinchoninic acid assay using a kit (Thermofisher, USA). Protein lysates were separated on an SDS-PAGE (12%) and then transferred to a PVDF membrane (Bio-Rad). The membrane was blocked by dipping in 3% skimmed milk (Bio-Rad) in PBS containing 0.1% Tween 20 (Amresco) (1xPBST) for 1.5 h with slow rocking at room temperature. IAV-associated matrix protein (M1) was detected by an anti-M1 antibody (1:500) (GTX-127356) as primary, and an HRP conjugated goat anti-rabbit IgG H&L (1:10,000) was used as a secondary antibody. The same blots were used after stripping for estimating β-actin as a loading control using an HRP-conjugated primary antibody against β-actin (1:10,000).

#### Immunofluorescence Imaging

A549 cells were plated (0.05 X 10^5^ cells/well) in a 24-well tissue culture plate containing sterile coverslips preplaced before seeding. Virus (Influenza-A/H_1_N_1_/WSN) with different dilutions of peptide was made in Serum-free media (SFM) containing TPCK-treated trypsin (2.5 µg/ml). To infect, cells were washed with 1XPBS (pH 7.4), then different dilutions were added to each well and incubated for 45-50 minutes at 37°C, 5% CO_2_. After removing the infection media, cells were washed twice with 1XPBS, and growth media containing 0.1% FBS, 0.3% BSA, and trypsin (2.5 μg/ml) was added and incubated for 15 hours at 37°C, 5% CO_2_. For immuno-staining, cells were first fixed using 4% paraformaldehyde, pH 7.4 (SIGMA) made in 1XPBS and then permeabilized for 15 minutes with 0.1% Triton X-100 (SRL) in 1XPBS at RT. Cells were washed with 1XPBS, and blocking solution (2% BSA in 1XPBS) was added for 1 hour at RT. The primary antibody, mouse monoclonal Anti M1 antibody (GTX-127356), was added in a 1:500 dilution for 90 minutes at RT. After that, the cells were washed five times with 1xPBS. Following this, cells were incubated with an Alexa Fluor®488 conjugated Secondary antibody (ab150113 from Abcam) diluted in 0.1% BSA in PBS (1:10000) for 1 hour in the dark. Cells were washed five times with PBST and mounted on glass slides using mounting media (fluoroShield with DAPI) (SIGMA). The slides were imaged using a confocal laser-scanning microscope (OLYMPUS).

#### Chikungunya virus infection inhibition assay

BHK-21 cells in 96-well plates were treated with different peptide concentrations for two hours, followed by infection with CHIKV-GFP (181/25) at 0.1 MOI for one hour. The inoculum was replaced by medium containing peptides and incubated for 24 hours. The cells were trypsinized and fixed in 4% buffered formaldehyde. The GFP-positive cells were assessed for virus infection using a flow cytometer (BD FACS Lyric), and the number of infected cells was determined using flow cytometry. After normalization with cell controls, the % infected cells in the peptide-treated cells were calculated after setting infected cells in the virus control as 100%.

**Figure S1:**
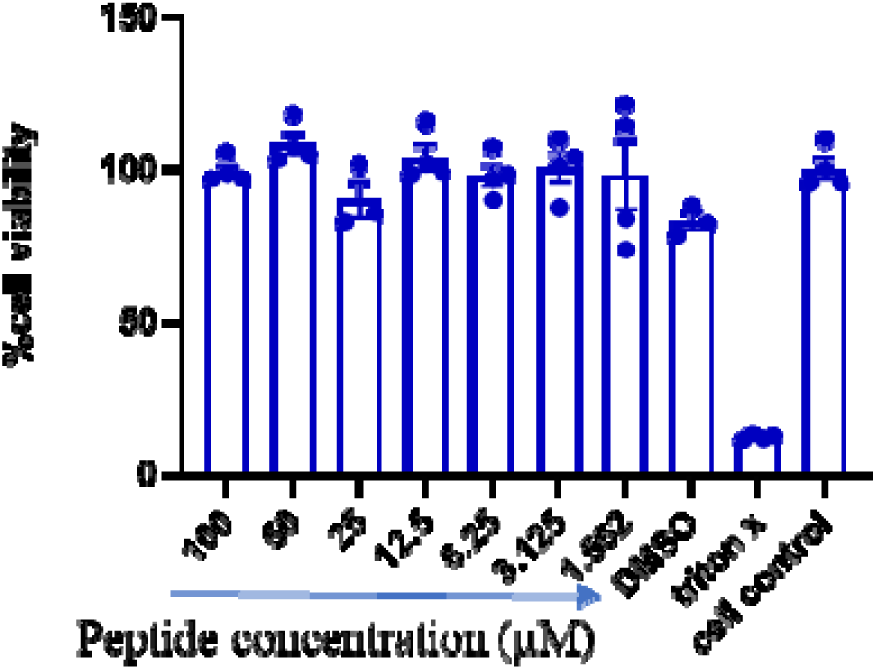
Cell viability in the presence of mGG21 by MTT assay. A549 cells were incubated with increasing concentrations of mGG21. Cell viability was assessed after 48 hours. Data shown are mean ± SEM of four replicates.

